# Real-time fluorescence and deformability cytometry — flow cytometry goes mechanics

**DOI:** 10.1101/187435

**Authors:** Philipp Rosendahl, Katarzyna Plak, Angela Jacobi, Martin Kraeter, Nicole Toepfner, Oliver Otto, Christoph Herold, Maria Winzi, Maik Herbig, Yan Ge, Salvatore Girardo, Katrin Wagner, Buzz Baum, Jochen Guck

**Affiliations:** Biotechnology Center, Center for Molecular and Cellular Bioengineering, Technische Universität Dresden, Dresden, Germany.; MRC Laboratory for Molecular and Cellular Biology, University College London, Gower Street, London, WC1E6BT, UK; Medical Clinic I, University Hospital Carl Gustav Carus, Technische Universität Dresden, Dresden, Germany; Microstructure Facility, Center for Molecular and Cellular Bioengineering, Technische Universität Dresden, Dresden, Germany

## Abstract

Cell mechanical characterization has recently approached the throughput of conventional flow cytometers. However, this very sensitive, label-free approach still lacks the specificity of molecular markers. Here we combine real-time 1D-imaging fluorescence and deformability cytometry (RT-FDC) to merge the two worlds in one instrument — opening many new research avenues. We demonstrate its utility using sub-cellular fluorescence localization to identify mitotic cells and test for their mechanical changes in an RNAi screen.

---

Flow cytometry (FCM) is the gold standard for single cell characterization in biological research and numerous clinical applications^1,2^. As cells are flowed through the cytometer’s cuvette, they are illuminated by a laser source, while detectors collect emitted fluorescence, as well as forward- and side-scattered light^2^. The analysis of scattered light provides information about cell size and shape in the absence of a molecular label.

Recently, mechanics has emerged as an orthogonal label-free way to obtain information at the cellular scale. Importantly, a cell’s mechanical state is tightly regulated in a cell-type and cell-state specific manner. Cell compliance therefore functions as a quantitative read-out of the state of the cytoskeleton^3-6^, which is influenced by progression through the cell cycle^7^, differentiation^8^, as well as patho-physiological processes, such as malignant transformation^9,10^ and by immune challenge during infections^11^. Due to the recent advent of several microfluidic techniques with a massive increase in throughput (i.e., up to thousands of cells/sec), mechanical phenotyping can now be performed at measurement rates approaching those of conventional flow cytometers^12-16^. Amongst many other applications, this opens up the possibility of carrying out large-scale screens for the genes that regulate cell mechanics — something that was previously limited by the low throughput (i.e., 10 – 100 cells/h) of techniques such as atomic force microscopy (AFM).

Label-free approaches provide quantitative information about cells but lack molecular specificity. Such molecular-level information can, however, be acquired in parallel by combining these approaches with fluorescent probes^2^. In FCM the intracellular distribution of the fluorescent signal can be obtained by fluorescence pulse shape analysis^17,18^. Slit-scan flow cytometers detect fluorescence in a thin light sheet and achieve 1-D fluorescence imaging with improved resolution^19,20^. Line scan cameras^21^ and advanced microscopy techniques based on non-linear optics^22,23^ even implement 2-D imaging. Thus, mechanical phenotyping, which is very sensitive to changes in cell state, is an ideal complement to the use of fluorescent molecular probes. The combination of both approaches promises to open up entirely new possibilities for scientific exploration.

Here, we present real-time fluorescence and deformability cytometry (RT-FDC), using a machine that combines the capabilities of 1D-resolved, fluorescence-based flow cytometry with mechanical phenotyping for the continuous analysis of large cell populations (**Fig. 1a**). The principle of operation is based on real-time deformability cytometry (RT-DC)^14^. Up to 100 cells/s are flowed through a microfluidic chip (**Supplementary Fig. 1**) and deformed without physical contact by hydrodynamic interactions in a constriction zone^24,25^ (**Fig. 1b**). Stroboscopic high-speed bright-field microscopy with real-time image processing captures and evaluates the cell contour. Simultaneously, three diode lasers excite fluorescence in a light sheet perpendicular to the channel axis (**Fig. 1a**, **Supplementary Fig 2a,b**) and avalanche photodiode detectors measure emitted fluorescence intensity in this region. If there is an object detected in the bright-field image, the custom algorithm extracts cross-sectional area and deformation (**Fig. 1b**), which can be related to the elastic modulus of the object^24,25^, and analyzes the respective fluorescence detector signal for peak maximum, width, and area.

**Figure 1.**
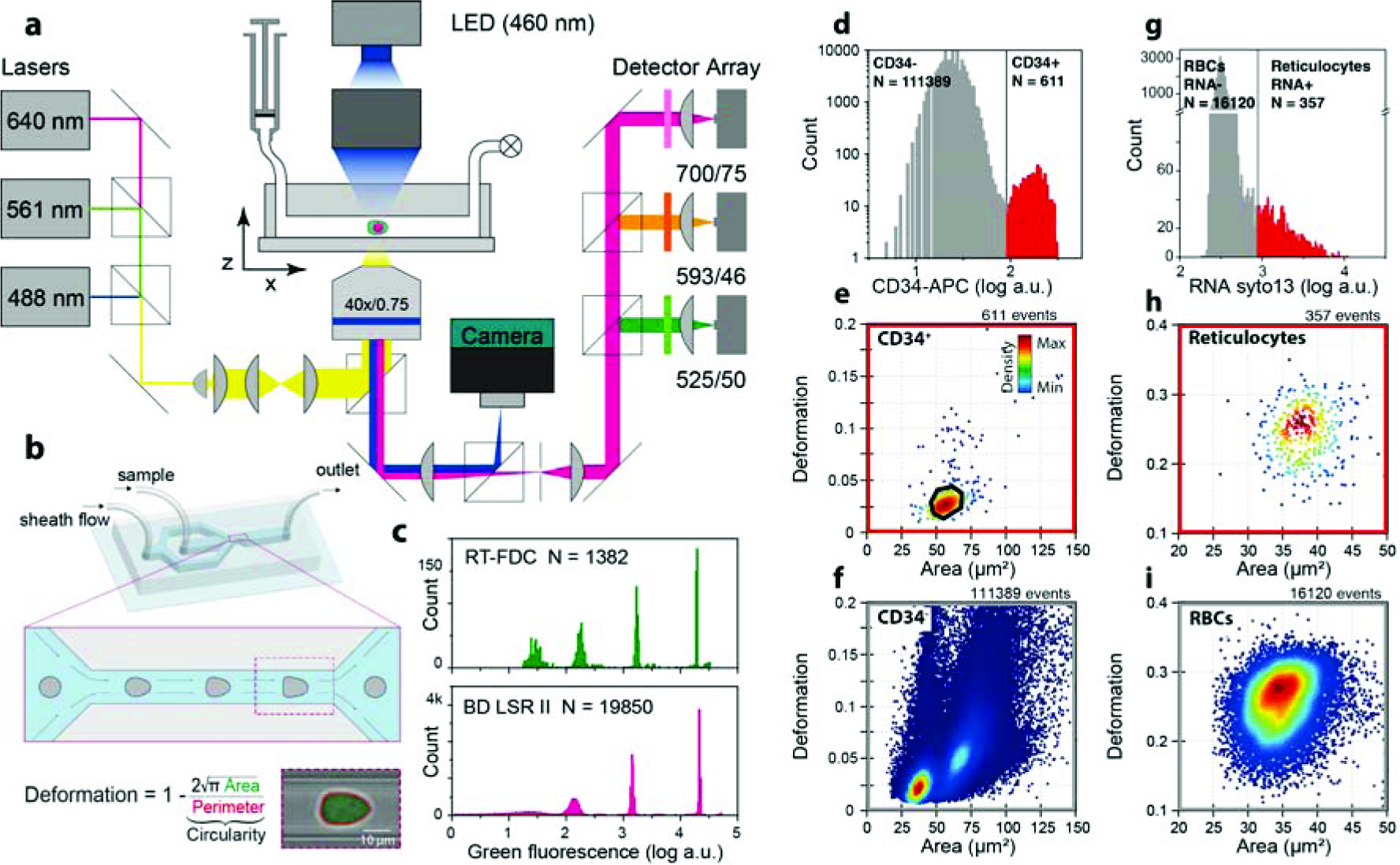
Real-time fluorescence and deformability cytometry. (**a**) Experimental setup. Cells are pumped through a microfluidic channel and imaged by bright-field microscopy, while three lasers excite, and avalanche photodiodes measure fluorescence. (**b**) In the narrow constriction zone cells deform by hydrodynamic interaction and deformation is determined by image processing. (**c**) Fluorescence intensity histogram of calibration beads as measured by RT-FDC (green) shows four populations, confirmed by state-of-the-art FCM (pink). (**d-i**) Representative plots from experiments for identification of subpopulations in mechanical fingerprints with fluorescent probes. (**d**) HSPCs are surface labeled with anti-CD34 antibodies conjugated to allophycocyanin (APC). Log-histogram of CD34-APC fluorescence, and gate used for classification. (**e**) CD34^+^ cells form a homogeneous population. Color of the dots indicates a linear density scale. Black polygon drawn around cluster corresponds to a gate used for calculating confusion matrix. (**f**) CD34^−^ cells show more spread in the deformation vs. area plot. (**g**) Log-histogram and gate for reticulocytes labeled with syto13. Mechanical fingerprint of (**h**) RNA-positive reticulocytes and (**i**) RNA-negative RBCs.

To evaluate RT-FDC as a technique, we first compared how fluorescence detection performs relative to standard FCM. The histograms in **Fig. 1c** show the intensities (fluorescence peak height) measured with RT-FDC and with a commercial flow cytometer (LSR II, Becton Dickinson) for calibration beads (see also **Supplementary Fig. 2c**). Both methods consistently identify four populations of objects with intensities over a dynamic range of three orders of magnitude. The synchronicity of the different fluorescence channels was checked using custom-made 2-color fluorescent agarose beads (see Methods, **Supplementary Fig. 2d**). To test the utility of RT-FDC — the combined fluorescence and deformability analysis — for biological and biomedical research, we then performed several series of experiments where we assessed the mechanical properties of subpopulations of fluorescent cells in a heterogeneous mixture without prior sorting.

**Figure 2.**
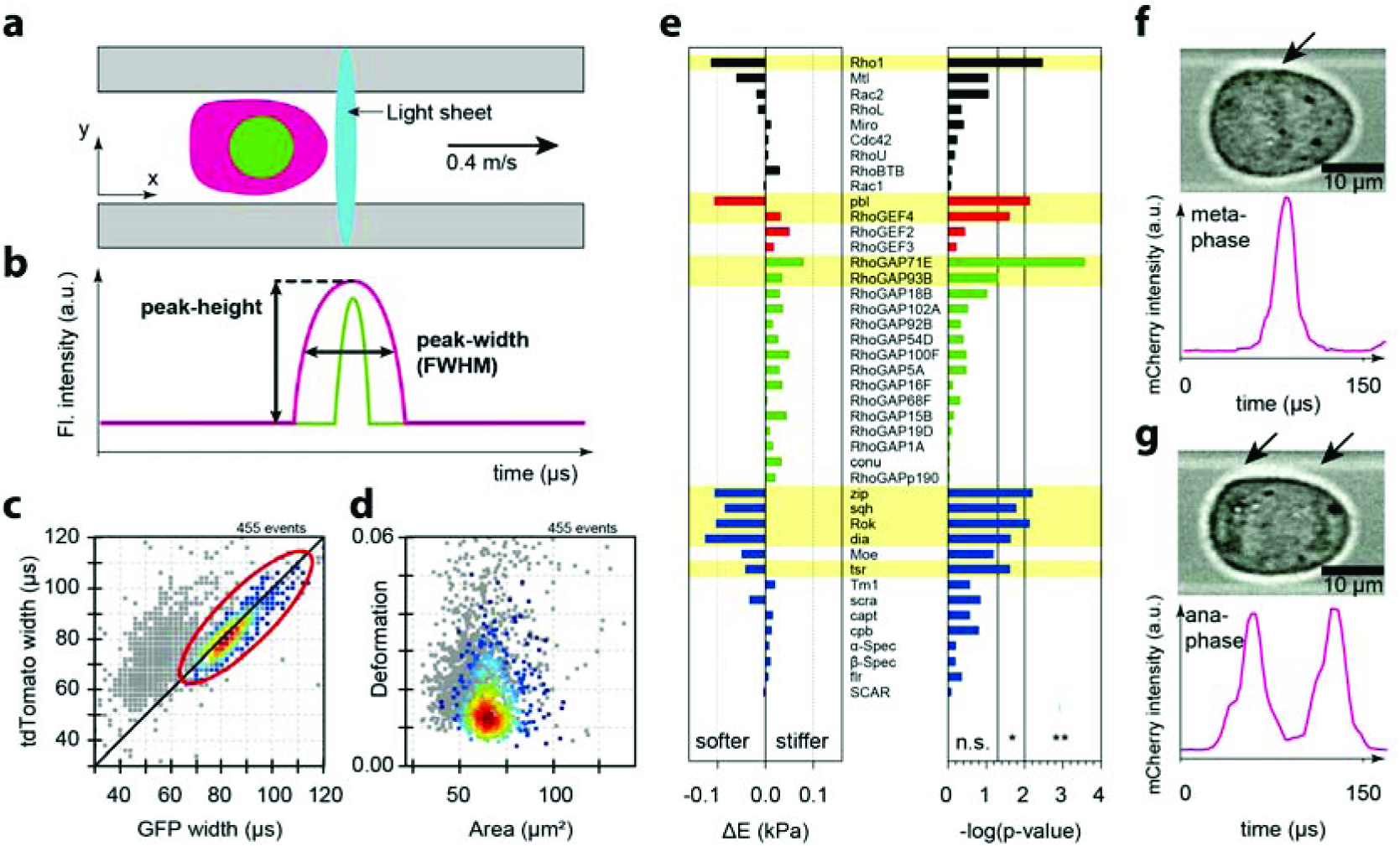
1D fluorescence imaging for identification of mitotic cells. (**a**) In the interrogation volume cells cross a 3 μm thick light sheet for fluorescence excitation. (**b**) Peak height and width analysis reveal sub-cellular fluorophore localization. (**c**) Nuclear envelope breakdown leads to equal nls-GFP and tdTomato peak widths. Mitotic cells along the diagonal can be gated with red polygon (cf. **Supplementary Fig. 5**). Interphase cells shown in grey. (**d**) Deformation vs. area diagram of mitotic sub-population. (**e**) Results of RNAi screen in Kc167 cell line for regulators of cell mechanics during mitosis. Change in elastic modulus and significance from linear mixed model evaluation are shown; hits are highlighted in yellow. (**f,g**) Sub-cellular localization of mCherry-H2B in HeLa cells. Cells in (**f**) metaphase and (**g**) anaphase exhibit one or two fluorescence peaks corresponding to one metaphase plate (arrow) or two separated chromatids (arrows), respectively.

Surface marker labeling is the standard approach for identification of many different cell types including hematopoietic stem and progenitor cells (HSPCs)^14^. Mechanical properties of HSPCs have been shown to be linked to their circulation and migration ability^8,26^ — essential aspects of successful HSCP homing when used for transplantation after chemotherapy. For validation of cell mechanics as a phenotypic marker for CD34^+^ HSPCs, granulocyte colony-stimulating factor (G-CSF) mobilized peripheral blood of human donors was measured (see Methods). After classification into CD34^+^ and CD34^−^ by means of fluorescence intensity using RT-FDC (**Fig. 1d**), the mechanical fingerprint (deformation vs. area plot, **Fig. 1e,f**) revealed that CD34^+^ cells had a mean size of 61.1 ± 1.3 μm^2^ and low deformation ranging from 0.03 to 0.05, corresponding to an elastic modulus of approximately 800 Pa, as confirmed with three different donors. CD34^−^ were either smaller (lymphoid cells)^14,27^ or larger (monocytes and granulocytes)^14,27^ and displayed a wider distribution in deformation. To test the potential power of sorting by means of deformation and cell-size alone — and without relying on fluorescence — a gate, as shown as black polygon in **Fig. 1e**, was set for validation of the mechanical phenotype as a marker. Assuming the fluorescence read-out as ground truth, classification by means of the mechanical fingerprint yielded a sensitivity of 76.5 % and specificity of 89.7 % (see confusion matrix **Supplementary Table 1**). The prevalence was 0.55 %, as confirmed with FCM (see **Supplementary Fig. 3**). These values would permit the strong enrichment of HSPCs in G-CSF mobilized blood, based on their morphological and mechanical phenotype alone. Fluorescence labeling could still be used for confirmation of successful sorting and enrichment on a small fraction of cells while the majority of unstained sample could be used for patient transplantation.

As a second test case, to demonstrate the applicability of RT-FDC for cell-permeable fluorescent dyes, we used RT-FDC to determine the mechanical properties of immature versus mature red blood cells (RBCs). Reticulocytes are the immature fraction of RBCs and represent normally about 5 – 15 ‰ of the total number of RBCs in circulating human blood^28^. During the first two days after being released to the blood stream, reticulocytes lose their cytoplasmic RNA as they mature to RBCs. This enabled us to use nucleic acid staining with syto13 to identify RNA-positive reticulocytes by fluorescence, as shown in the histogram **Fig. 1g** and to determine the mechanical profile of both cell types (**Fig. 1h-i**). Reticulocyte numbers relative to RBCs were in agreement with a state-of-the-art automatic blood count (**Supplementary Fig. 4a**). In addition, in line with previous reports on reticulocytes^29^, which had been restricted to labor-intensive micropipette and micro-pore assays with limited number of cells, we found human reticulocytes in whole blood samples to be slightly larger and less deformed compared to RBCs (**Fig. 1h-i**, **Supplementary Fig. 4b, Supplementary Table 2**). Increased reticulocytosis has been linked with increased morbidity in infants with sickle cell anemia^30^ and RBC distribution width is also used as a predictor of poor clinical outcome in settings of various diseases including coronary heart disease^31^. Adding RT-FDC phenotyping to the reticulocyte indices in clinical use could therefore provide a means of monitoring changes of blood rheology in haematological diseases^27,31^.

In addition to detecting merely the fluorescence intensity of a cell, the setup also permits 1D fluorescence imaging. As the light sheet has a thickness of 3 μm (**Fig. 2a**, **Supplementary Fig. 2a,b**), which is smaller than most eukaryotic cells, and cells are traveling through the channel with constant speed, the temporal shape of a fluorescence peak depends on the subcellular distribution of the fluorophore along the channel direction (**Fig. 2b**). This enables RT-FDC to provide spatial information about the localization of the fluorescence signal within cells. Taking advantage of this feature, we used 1D imaging to detect the nuclear localization of a fluorescent protein by measuring the difference of peak width for cells expressing nuclear localization sequence (nls) tagged and cytoplasmic (cyt) localized fluorescent proteins (**Supplementary Fig. 5**). Since, in animal cells, passage through the cell cycle involves changes in nuclear permeability, we were then able to exploit this technology to selectivity analyze the mechanical profiles of mitotic cells. In order to develop a system in which we could screen for changes in mitotic cell mechanics, we chose to use the *Drosophila* cell line Kc167. This enabled us to benefit from the lack of genetic redundancy in the fly genome, relative to mammalian cells, and the efficacy of RNAi in fly cell culture. In addition, these cells grow well at room temperature, facilitating the analysis^32^.

Mitotic cells were detected in a heterogeneous population based on nuclear envelope breakdown, when nls-GFP leaked into the cytoplasm and GFP peak width was identical to cytosolic tdTomato peak width (**Fig. 2c**), and their specific mechanical fingerprint was analyzed (**Fig. 2d**). We then screened for genes that regulate mitotic cell mechanics. We investigated the effects of 42 gene depletions — chosen to target known regulators of the cortical actin cytoskeleton. Each gene was targeted by two independent dsRNAs, in three biological replicates and two different flow rates — a total of over 500 individual measurements. To compensate for potential cell size changes due to gene depletions, we utilized a numerical model^25^ for calculating elastic modulus for cells deformed in RT-FDC. A linear mixed model^33^ was then used to compare treatment and control samples (results for analysis of elastic modulus changes are summarized in **Fig. 2e**). The detailed analysis pipeline and all screening results can be found in **Supplementary Fig. 6**, and **Supplementary Table 3**.

This screen identified a set of known and novel genes that regulate mitotic cortical mechanics. Importantly, as expected, we were able to detect a loss of cortical stiffness that resulted from disrupting the Rho signaling cascade. These included RhoGEF (Pbl/Ect2); Rho itself; Rock, a downstream kinase; Myosin (Zipper/Squash), and a downstream actin nucleator (Dia). Importantly, these results are in good agreement with data obtained by other methods for measuring cell mechanics of mitotic cells^34,35^. Further, the screen showed that less-well studied RhoGAP proteins are likely to counter the RhoGEFs in the control of mitotic cell mechanics. Thus, using RT-FDC it is possible to screen for regulators of cell mechanics even in sub-populations of cells. A detailed discussion of the screen results can be found in the **Supplementary Discussion**.

After showing the ability of the setup to detect differences between nuclear and cytoplasmic compartments we tested whether we are able to detect intracellular structures smaller than the entire nucleus such as separating chromatids during mitosis. In **Fig. 2f,g** we show two examples of mitotic HeLa cells with fluorescently labeled histone 2B for tracking chromosomes. We were able to detect cells with single and double peaks of mCherry fluorescence signal, which we associated with metaphase and anaphase cells, respectively (condensed chromosomes are also visible in bright field images of cells). An important aspect for the detection of separated chromatids is the proper alignment of cells elongated along the separation axis with the channel axis, which is ensured by the hydrodynamic forces acting on the elongated cells. Thus, it is possible to detect mitotic cells and sub-stages of mitosis. Especially for mechanical studies of short cell cycle phases such as anaphase the high-throughput capabilities of RT-FDC will be crucial to detect the very small fractions in unsynchronized samples.

In summary, these experiments demonstrate that RT-FDC detects the most common fluorescent probes: surface antibody labels, cell permeable dyes, and fluorescent proteins. RT-FDC combines the molecular specificity of fluorescent probes with the sensitivity of the functional marker cell mechanics, enabling researchers to find correlations between both, or gate for rare cells in large samples for mechanical analysis. Cell mechanics was evaluated as a label-free marker for HSPCs against CD34-fluorescence, showing that it should be possible to enrich for cells using mechanics alone. In the future, this may aid or even substitute the current label-based purification of HSCPs. We also showed how RT-FDC can be used to reveal an otherwise invisible subpopulation of reticulocytes amongst the RBCs, and to assess their relative mechanical properties. Finally, our results show that the technique is an ideal method to screen for regulators of cell mechanics. Screens focused on mechanical properties will benefit from the high throughput per sample (100 cells/s) but also from the short time needed to complete one experiment (15 min) permitting more than 500 single experiments performed for the presented screen on candidate genes for regulation of cell mechanics during mitosis. Future investigations of mitotic cell mechanics might take into account information encoded in the fluorescence peak with greater detail, as demonstrated for the H2B HeLa cells for resolving meta-, ana- and telophase. This aspect highlights the need for simultaneous fluorescence and mechanical read-out: If cells would have to be presorted by FACS before mechanical measurements by any given method, time sensitive phases, such as mitosis, could not be resolved due to experimental delay. Only simultaneous read-out retrieves the full information on a single cell level, going beyond the state of the art of what is achievable using flow cytometry or deformability cytometry alone.

## Online Methods

### Experimental Setup

**Figure 1** shows the experimental setup used throughout this study. The microfluidic chip, synchronized pulsed LED illumination (AcCellerator L1, Zellmechanik Dresden), syringe pump (Nemesys, cetoni) and the bright field imaging (MC1365, Mikrotron) are adapted from RT-DC.^14^ For fluorescence excitation, three solid state lasers (OBIS 488 nm LS 60 mW; OBIS 561 nm LS 50 mW; OBIS 640 nm LX 40 mW, Coherent Deutschland) in combination with adjustable mounted dichroic mirrors (561-594R; 473-491R; 1064R, Semrock) create a combined beam. This beam is expanded and collimated by two achromatic lenses (AC080-010-A-ML; AC254-040-A-ML, Thorlabs) and then focused to a sheet in the image plane of the microscope with a cylindrical lens (LJ1695RM-A, Thorlabs). Excitation light is coupled into the microscope (Axio Observer.Z1, Carl Zeiss Microscopy) through the modified backport for epifluorescence illumination. Below the 40x objective (EC Plan-NEOFLUAR 40x/0.75, Carl Zeiss Microscopy) a QuadLine beamsplitter (zt405/488/561/640rpc, Chroma Technology) reflects excitation light towards the specimen but transmits light emitted by the sample to the detector assembly. A second beam splitter (zt 473 RDCXT, Chroma Technology) separates light from the LED illumination with 460 nm to the CMOS camera (MC1365, Mikrotron) for bright field imaging and light of higher wave lengths to an adjustable slit (VA100C/M, Thorlabs) that is in the image plane of the microscope and part of the confocal detector assembly. The collimated beam is separated into three fluorescence channels: FL-1 (FF555-Di03; FF03-525/50, Semrock), FL-2 (zt 633 RDC, Chroma Technology Corp; FF01-593/46, Semrock) and FL3 (700/75 ET, Chroma Technology Corp) and finally focused (LA1951-A-ML, Thorlabs) on the three avalanche photodiode detectors (MiniSM10035, SensL Corporate). The analog detector signals are digitized by a PCIe card (NI PCIe-6531, National Instruments Germany) at a sample rate of 1 Msamples/s (shared between channels) and stored in a circular buffer of 1 Msamples for real-time analysis. By deriving the trigger signal for image acquisition from the same clock as the detector data acquisition, the image and fluorescence acquisition are synchronized. If an object is detected in an image, the corresponding part of the detector signal is fetched from the circular buffer and also analyzed. For image processing the OpenCV 3.1 computer vision library (http://opencv.org) was used wherever possible. Image processing includes the following steps: subtraction of a background image (rolling average of last 100 camera images), threshold operation, finding contours in binary image by a border following algorithm^30^, determining particle cross sectional area, deformation (1 - circularity) and position of the contour. In case these parameters match the gates set before, the corresponding fluorescence detector signal is fetched from the circular buffer and analyzed for the maximum value, the peak width (FWHM), the peak area and the peak position. These parameters are stored together with results from image processing, an image of the cell and the fluorescence peak to the disk of the computer for later analysis and instantly plotted on the screen.

For characterization of the aperture function (sensitivity as a function of position)^31^, fluorescent beads with a size of 3 μm (BD FACSDiva CS&T Research Beads, Becton Dickinson) were embedded in 1 % agarose gel (low gelling temperature A0701-100G, Sigma-Aldrich Co. LLC) on a microscope slide (Thickness 2, GlaswarenfabriknKarl Hecht) and moved through the detection volume using the automated microscope stage while recording the detector signals in order to obtain the combined fluorescence excitation and emission efficiency for a field of 30 μm x 60 μm as shown in **Supplementary Figure 2a**. Since the microscope slide was the same used for the microfluidic chips and the refractive index of 1 % agarose gel is very close to that of water (*n =* 1.34), it is assumed that the light sheet in the microfluidic channel and the bead test slide have a very similar shape. The resulting thickness of the light sheet (FWHM) along the channel axis is presented in **Supplementary Figure 2b** and is about 3 μm for all fluorescence channels. This is also the size of the fluorescent bead and underlines the capability of the device to detect localization of fluorophores in cell compartments such as the nucleus that have a size of about 5 μm.

The dynamic range of the fluorescence detection system was tested with flow cytometer calibration beads (Rainbow Calibration Particles RCP-35-5, Spherotech). As expected, four (FL-1) or three (FL-2; FL-3) populations show up in the histogram of the fluorescence peak heights shown in **Supplementary Fig. 2c** in agreement with reference measurements provided in the manufacturer’s manual and own measurements performed on a LSR II flow cytometer (Becton Dickinson). Fluorescence peak width detection in different channels is compared in **Supplementary Fig. 2d**, which shows data from 2-color fluorescent agarose beads with heterogeneous sizes. These beads were produced from a mixture of two ultra-low gelling point agarose solutions, one functionalized with Alexa488, the other with Alexa633. Crosslinking was achieved by decreasing temperature below the gelling point resulting in stable beads that can be stored for weeks. The linear fit shows that the peak widths in the green and the read channel (FL-1 and FL-3) are proportional with a slope close to one and a small offset of - 0.1 μs, demonstrating that peak width measurements can be compared across channels to study fluorophore localization. The particle speed in the flow for this measurement is about 0.4 m/s so the particle diameters range from 5 μm to 35 μm based on the fluorescence signal peak widths. All the setup characterization experiments were performed in three technical repeats.

### Experimental Procedure

For all experiments cells or micro particles were suspended in phosphate buffered saline without magnesium/calcium (PBS^-^) complemented with 0.5 % or 0.6 % (w/v) methylcellulose (Alfa Aesar) to slow down sedimentation during the experiment. The methylcellulose solution was calibrated with a falling ball viscometer (Haake Typ C, Thermo Fisher Scientific) to a viscosity of 15.0 ± 0.8 mPa·s and 25 ± 0.6 mPa·s at 24 °C for 0.5 % and 0.6 % methylcellulose respectively. Prior to the experiments the cell suspension was aspirated into PEEK tubing (Upchurch, Thermo Fisher Scientific) connected to a 1 mL plastic syringe (Becton Dickinson) at a flow rate of 1 μL/s avoiding high stress on the specimen. Then the tubing was connected to the microfluidic chip (shown in **Fig. 1b** **and Supplementary Fig. 1**) were constant flow was generated by combination of sheath and sample flow with a ratio of 3:1. After equilibration of the flow for two minutes, the measurement was started. The camera’s region of interest (256 pixels x 96 pixels) was positioned at the end of the 300 μm long channel region of the chip to ensure cell deformation had reached a steady state. Laser powers can be adjusted in a range between 1 mW and 60 mW to match the detector’s dynamic range in an optimal way. Due to low spectral overlap between fluorophores used in our experiments no compensation was applied to raw fluorescence data. Compensation procedure may however be performed following standard methods. Depending on the cell concentration in the sample, measurements ran for one to a few minutes and obtained typically 2,000 - 100,000 events at rates of up to 100 events per second. Higher throughput rates were not yet possible because multiple cells in the image might impede correct assignment of fluorescence peaks to cells in the image data. Post experimental data analysis was performed with the following programs: ShapeOut (Zellmechanik Dresden, https://github.com/ZELLMECHANIK-DRESDEN/ShapeOut) for plotting and gating, while OriginPro 9.0 (OriginLab Corporation) was used for more detailed data inspection, plotting, and fitting.

### Hematopoietic stem cell isolation and preparation

With approval for the study (EK221102004) from the ethics committee of the Technische Universität Dresden, we analyzed the apheresis product from G-CSF mobilized peripheral blood of three healthy human donors with their informed consent in accordance with the guidelines of good practice and the Declaration of Helsinki. Before measuring with RT-FDC, HSPCs were stained for 30 min with a CD34-APC conjugated antibody (#555824; Becton Dickinson), then pelleted by centrifugation (200 g, 5 min), and resuspended in 0.6 % methylcellulose solution.

### Reticulocyte and red blood cell preparation

With approval for the study (EK89032013) from the ethics committee of the Technische Universität Dresden, we obtained blood from three healthy donors with their informed consent in accordance with the guidelines of good practice and the Declaration of Helsinki. Capillary blood was collected after finger prick with a 21G, 1.8 mm safety-lancet (Sarstedt) For sample preparation 2 μl of blood was diluted in 1 mL of 0.5 % methylcellulose solution complemented with 2.5 μM syto13 nucleic acid stain (Thermo Fisher Scientific). Measurements for three healthy donors were conducted after 5 minutes of incubation and took typically 10 minutes. To validate stability of mechanical phenotype and fluorescence staining in methylcellulose measurements, a 1 hour time course with measurements every 10 minutes was performed for all donors. To validate counting results, blood from venipuncture with 21 gauge needle (Multifly, Sarstedt) was collected in EDTA S (S-Monovette 1.4 ml 9NC, Sarstedt) for measurements with RT-FDC and parallel full blood count on a state-of-the-art device (XE-5000, Sysmex Deutschland) at the Institute for Clinical Chemistry and Laboratory Medicine at the Carl Gustav Carus University Hospital at the TU Dresden. The experiment and analysis were performed for three healthy donors.

### Cell lines and culture

The stable cell line of H2B mCherry HeLa was a kind gift of the lab of Matthieu Piel (Institute Curie, Paris). All HeLa cells were grown in standard DMEM media supplemented with 10 % fetal bovine serum and 2 mM L-Glutamine. Transfection of HeLa cells was performed using Effectene reagent (Quiagen) according to manufacturer’s directions. Cells were grown for 48 h after transfection and dead cells were removed by washing away loosely attached cells. Prior to RT-FDC measurements adherent cells were trypsinized, collected by centrifugation, and resuspended in 0.5 % methylcellulose buffer.

For mitotic synchronization of H2B mCherry HeLa line cells were allowed to grow to 35% confluency and then synchronized with 100 nM Nocodazole for 5 h. Mitotic cells were collected by shake-off and Nocodazole was washed away. Cells were then allowed to progress through mitosis in full media for 15 min (metaphase cells) or 1 h (anaphase cells).

Kc167 *Drosophila* cells were grown in M3 Shields and Sang medium supplemented with 10 % FBS. For detection of mitotic cells, we engineered a stable Kc167 cell line that expresses tdTomato cytosolic protein and NLS(GFP)2(GST), a protein that stably localizes to the nucleus during interphase. At the entry to mitosis, the nuclear envelope breaks down and the nuclear GFP signal is redistributed to the cytoplasm where it co-localizes with tdTomato. With analysis of fluorescence peak widths, interphase cells (narrow GFP peak and broad tdTomato peak) were distinguished from mitotic cells (both peaks of the same width; see **Fig. 2c**). Due to progression through mitosis and cytokinesis, cells in late mitotic stages obtain non-spherical shapes due to elongation and furrow constriction before division (anaphase and telophase of mitosis), which may bias the deformation readout of RT-FDC. To avoid this effect for the following experiments, we focused on prometaphase cells by synchronization with colchicine, a drug that inhibits formation of the mitotic spindle. Cells were synchronized with 4 μM colchicine for 5 h for enrichment of the mitotic cell fraction. RNA interference experiments were conducted as described before^32^. The treatment with colchicine lead to an increase in mitotic cell size; it did, however, not affect the mechanical phenotype of mitotic cells (**Supplementary Table 4**) and, as expected, it led to enrichment of the mitotic cell fraction (**Supplementary Fig. 7a,b**).

### RNA interference screening

For each of the 42 candidate genes (**Supplementary Table 5**) RNA interference was performed with two non-overlapping double stranded RNA sequences (dsRNA). Data from three independent experiments performed on different days was collected for each RNA sequence and compared to negative controls acquired on the same day (see **Supplementary Fig. 6**). For evaluation of the measured elastic modulus values, a linear mixed model^33^ comparison was performed with the software ShapeOut (Zellmechanik Dresden), for which data for each gene (different days, different sequences) was pooled. We defined rejection of the null-model with a p-value smaller than 0.05 as a hit. Only if the effect was visible for both flow rates (0.04 μL/s and 0.06 μL/s) the gene was considered to play a role in regulation of mitotic cell mechanical properties.

### Data availability

The raw RT-FDC data is available upon request as TDMS files that can be read, processed and analyzed by “ShapeOut”, an open source software that can be found on GitHub (https://github.com/ZELLMECHANIK-DRESDEN/ShapeOut). Polygon filters used for selecting the populations of cells of interest are available upon request and are also compatible with “ShapeOut” software.

### Statistical analysis

Statistical analysis was performed using the linear mixed model method (LMM) integrated into “ShapeOut” software. The method is described in detail in^33^.

### Code availability

Source code for the custom C++ acquisition software is made available upon request.

### Materials and Correspondence

Requests regarding material and general request shall be addressed to Jochen Guck.

## Author contributions

J.G. conceived the project. P.R. built the experimental setup. O.O. and C.H. supported the optics design. P.R. programmed the real-time data evaluation software. K.P. designed and performed experiments on KC167 and HeLa cells and analyzed the data with B.B.’s support. P.R. and N.T. performed experiments with RBCs. A.J., M.K., and P.R. performed experiments on HSPCs and analyzed the data. P.R. performed and analyzed experiments for device characterization. M.H., O.O., and Y.G. developed the linear mixed model algorithm for statistical evaluation. K.W. and S.G. produced fluorescent beads for device characterization. M.W. performed reference measurements with calibration beads on FCM. S.G. produced masters for microfluidic chips. J.G. and B.B. supervised the project. P.R., K.P. and J.G. wrote the manuscript. All authors reviewed the manuscript.

## Acknowledgements

We thank D. Klaue, M. Urbanska, F. Rosendahl, and I. Richter for helpful discussions and technical support. We thank M. Schürmann for refractive index measurements of agarose. We thank M. Piel for the donation of the HeLa H2B mCherry cell line.

Financial support from the Alexander-von-Humboldt Stiftung (Humboldt-Professorship to J.G.), Sächsisches Ministerium für Wissenschaft und Kunst (TG70 grant to O.O. and J.G.; European Fund for Regional Development — EFRE to S.G. and J.G.), an ERC Starting Grant (starting grant “LightTouch” #282060 to J.G.), the DFG Center for Regenerative Medicine of the Technische Universität Dresden (seed grant to J.G.), the DFG KFO249 (GU 612/2-2 grant to J.G.), a DKMS ‘Mechthild Harf Research Grant’ (DKMS-SLS-MHG-2016-02 to A.J.), and Cancer Research UK (C1529/A17343 grant to B.B.) is gratefully acknowledged.

## References

1. Picot, J., Guerin, C. L., Le Van Kim, C. & Boulanger, C. M. Flow cytometry: Retrospective, fundamentals and recent instrumentation. Cytotechnology 64, 109–130 (2012).

2. Shapiro, M. H. Practical Flow Cytometry. (John Wiley & Sons, Inc., 2003). doi:10.1002/0471722731

3. Elson, E. L. Cellular mechanics as an indicator of cytoskeletal structure and function. Annu. Rev. Biophys. Biophys. Chem. 17, 397–430 (1988).

4. Fletcher, D. & Mullins, R. Cell mechanics and the cytoskeleton. Nature 463, 485–492 (2010).

5. Xu, W. et al. Cell Stiffness Is a Biomarker of the Metastatic Potential of Ovarian Cancer Cells. PLoS One 7, e46609 (2012).

6. Kasza, K. E. et al. The cell as a material. Curr. Opin. Cell Biol. 19, 101–7 (2007).

7. Matthews, H. K. et al. Changes in Ect2 localization couple actomyosin-dependent cell shape changes to mitotic progression. Dev. Cell 23, 371–83 (2012).

8. Ekpenyong, A. E. et al. Viscoelastic properties of differentiating blood cells are fate-and function-dependent. PLoS One 7, e45237 (2012).

9. Guck, J. et al. Optical deformability as an inherent cell marker for testing malignant transformation and metastatic competence. Biophys J 88, 3689–3698 (2005).

10. Tse, H. T. K. et al. Quantitative diagnosis of malignant pleural effusions by single-cell mechanophenotyping. Sci. Transl. Med. 5, 212ra163 (2013).

11. Worthen, G. S., Schwab, B. I., Elson, E. L. & Downey, G. P. Mechanics of stimulated neutrophils: cell stiffening induces retention in capillaries. Science (80-.). 245, 183–186 (1989).

12. Gossett, D. R. et al. Hydrodynamic stretching of single cells for large population mechanical phenotyping. Proc. Natl. Acad. Sci. U. S. A. 109, 7630–5 (2012).

13. Dudani, J. S., Gossett, D. R., Tse, H. T. K. & Di Carlo, D. Pinched-flow hydrodynamic stretching of single-cells. Lab Chip 13, 3728 (2013).

14. Otto, O. et al. Real-time deformability cytometry: on-the-fly cell mechanical phenotyping. Nat. Methods 12, 199–202 (2015).

15. Byun, S. et al. Characterizing deformability and surface friction of cancer cells. Proc. Natl. Acad. Sci. U. S. A. 110, 7580–5 (2013).

16. Lange, J. R. et al. Microconstriction Arrays for High-Throughput Quantitative Measurements of Cell Mechanical Properties. Biophys. J. 109, 26–34 (2015).

17. Hoffman, R. A. in Current Protocols in Cytometry 1.23.1-1.23.17 (John Wiley & Sons, Inc., 2009). doi:10.1002/0471142956.cy0123s50

18. Ramdzan, Y. M. et al. Tracking protein aggregation and mislocalization in cells with flow cytometry. Nat. Methods 9, 467–470 (2012).

19. Johnston, R. G., Bartholdi, M. F., Hiebert, R. D., Parson, J. D. & Cram, L. S. Slit-scan flow cytometer for recording simultaneous waveforms. Rev. Sci. Instrum. 56, 691 (1985).

20. Gray, J. W., Peters, D., Merrill, J. T., Martin, R. & Van Dilla, M. a. Slit-scan flow cytometry of mammalian chromosomes. J. Histochem. Cytochem. 27, 441–444 (1979).

21. Basiji, D. A., Ortyn, W. E., Liang, L., Venkatachalam, V. & Morrissey, P. Cellular image analysis and imaging by flow cytometry. Clin. Lab. Med. 27, 653–670 (2007).

22. Lau, A. K. S. et al. Optofluidic time-stretch imaging — an emerging tool for high-throughput imaging flow cytometry. Lab Chip 16, 1743–1756 (2016).

23. Diebold, E. D., Buckley, B. W., Gossett, D. R. & Jalali, B. Digitally synthesized beat frequency multiplexing for sub-millisecond fluorescence microscopy. Nat Phot. 7, 806–810 (2013).

24. Mietke, A. et al. Extracting Cell Stiffness from Real-Time Deformability Cytometry: Theory and Experiment. Biophys. J. 109, 2023–2036 (2015).

25. Mokbel, M. et al. Numerical Simulation of Real-Time Deformability Cytometry To Extract Cell Mechanical Properties. ACS Biomater. Sci. Eng. acsbiomaterials.6b00558 (2017). doi:10.1021/acsbiomaterials.6b00558

26. Kräter, M. et al. Bone marrow niche-mimetics modulate HSPC function via integrin signaling. Sci. Rep. 7, 2549 (2017).

27. Toepfner, N. et al. Detection Of Human Disease Conditions By Single-Cell Morpho-Rheological Phenotyping Of Whole Blood. bioRxiv (2017).

28. d’Onofrio, G. et al. Simultaneous Measurement of Reticulocyte and Red Blood Cell Indices in Healthy Subjects and Patients With Microcytic and Macrocytic Anemia. Blood 85, 818–823 (1995).

29. Waugh, R. E. Reticulocyte rigidity and passage through endothelial-like pores. Blood 78, 3037–42 (1991).

30. Meier, E. R. et al. Increased Reticulocytosis during Infancy Is Associated with Increased Hospitalizations in Sickle Cell Anemia Patients during the First Three Years of Life. PLoS One 8, (2013).

31. Felker, G. M. et al. Red Cell Distribution Width as a Novel Prognostic Marker in Heart Failure: Data From the CHARM Program and the Duke Databank. J. Am. Coll. Cardiol. 50, 40–47 (2007).

32. Baum, B. & Cherbas, L. Drosophila cell lines as model systems and as an experimental tool. Methods Mol. Biol. 420, 391–424 (2008).

33. Herbig, M. et al. in Flow Cytometry Protocols (eds. Hawley, R. & Hawley, T.) (Humana Press, 2017). doi:10.1007/978-1-4939-7346-0

34. Chugh, P. et al. Actin cortex architecture regulates cell surface tension. Nat. Cell Biol. 19, 689–697 (2017).

35. Ramanathan, S. P. et al. Cdk1-dependent mitotic enrichment of cortical myosin II promotes cell rounding against confinement. Nat. Cell Biol. 17, 148–159 (2015).

